# Reconstitution of intestinal stem cell niche *in vitro* with pharmacological inhibitors or L-WRN conditioned medium differentially regulates epithelial proliferation, differentiation and barrier function in rabbit caecum organoids

**DOI:** 10.1101/2020.04.02.021550

**Authors:** Eloïse Mussard, Cécile Pouzet, Virginie Helies, Géraldine Pascal, Sandra Fourre, Claire Cherbuy, Aude Rubio, Nathalie Vergnolle, Sylvie Combes, Martin Beaumont

**Affiliations:** GenPhySE, Université de Toulouse, INRAE, ENVT, F-31326, Castanet Tolosan, France; Fédération de Recherche FR3450 (Agrobiosciences, Interactions et Biodiversité), Plateforme Imagerie-Microscopie, CNRS, Université Toulouse, 31326, Castanet Tolosan, France; GeT-PlaGe, Genotoul, INRAE, Castanet Tolosan, France; Micalis Institute, INRAE, AgroParisTech, Université Paris-Saclay, 78350, Jouy-en-Josas, France; IRSD, Université de Toulouse, INSERM, INRAE, ENVT, UPS, U1220, CHU Purpan, CS60039, 31024, Toulouse, France

**Keywords:** enteroid, spheroid, epithelium, culture medium, monolayer

## Abstract

Intestinal organoids are self-organized 3-dimensional (3D) structures formed by a single layer of polarized epithelial cells. This innovative *in vitro* model is highly relevant to study physiology of the intestinal epithelium and its role in nutrition and barrier function. However, this model has never been developed in rabbits, while it would have potential applications for biomedical and veterinary research. Here, we cultured rabbit caecum organoids with either pharmacological inhibitors (2Ki medium) or L-WRN cells conditioned medium (L-WRN CM) to reconstitute the intestinal stem cell niche *in vitro*. Large spherical organoids were obtained with the 2Ki medium and this morphology was associated with a high level of proliferation and stem cells markers gene expression. In contrast, organoids cultured with L-WRN CM had a smaller diameter; a greater cell height and part of them were not spherical. When the L-WRN CM was used at low concentration (5%) for two days, the gene expression of stem cells and proliferation markers were very low, while absorptive and secretory cells markers and antimicrobial peptides were elevated. Epithelial cells within organoids were polarized in 3D cultures with 2Ki medium or L-WRN CM (apical side towards the lumen). We cultured dissociated organoid cells in 2D monolayers, which allowed accessibility to the apical compartment. Under these conditions, actin stress fibers were observed with the 2Ki medium, while perijonctionnal localization of actin was observed with the L-WRN CM suggesting, in 2D cultures as well, a higher differentiation level in the presence of L-WRN CM. In conclusion, rabbit caecum organoids cultured with the 2Ki medium were more proliferative and less differentiated than organoids cultured with L-WRN CM. We propose that organoids cultured with the 2Ki medium could be used to rapidly generate *in vitro* a large number of rabbit intestinal epithelial stem cells while organoids cultured with the L-WRN CM represent a suitable model to study differentiated rabbit epithelium.

## Introduction

The intestinal epithelium is formed by a monolayer of cells located at the surface of the gut mucosa, facing the luminal environment (Peterson and Artis, 2014). Intestinal epithelial cells (IEC) play a pivotal role for both nutrition (digestion, nutrient absorption, hormone secretion) and barrier function (formation of tight junctions, secretion of mucus and antimicrobial peptides and detection of microbes). This pleiotropic function of the intestinal epithelium is linked to the presence of specialized differentiated IEC (absorptive, paneth, goblet and enteroendocrine cells) that arise from intestinal stem cells (ISC) located at the bottom of epithelial crypts (Barker, 2014). Recent progress in the characterization of the ISC niche (i.e. microenvironment) allowed the development of intestinal organoid culture *in vitro* (Sato et al., 2009). These self-organized 3D structures are formed by a closed lumen surrounded by a single layer of polarized IEC (Sato et al., 2009). Organoids closely replicate epithelial physiology and have been used as an efficient model to study nutrient transport, host-microbe interactions or pathological processes (Clevers, 2016; Lukovac et al., 2014; Zietek et al., 2015). They can also be used for pharmacological studies (Sébert et al., 2018).

Organoids are obtained by seeding tissue-derived ISC in an extracellular matrix (e.g. Matrigel) with growth factors reproducing *in vitro* the main ISC niche signaling pathways, i.e. high Wnt and low bone morphogenic protein (BMP) activation (Date and Sato, 2015; Gehart and Clevers, 2019; Holmberg et al., 2017). Thus, organoid culture medium usually contains three proteins: Noggin (inhibiting the BMP pathway), Wnt3a and R-spondin (activating the Wnt pathway) (Sato et al., 2009, 2011). Conditioned medium (CM) of engineered cell lines is a cost-effective and efficient source for these factors, when compared to purified recombinant proteins (Holmberg et al., 2017). For instance, the L-WRN cells are mouse L cells secreting Wnt3a, R-spondin 3 and Noggin which CM can be used to grow intestinal organoids (Miyoshi and Stappenbeck, 2013; VanDussen et al., 2015, 2019). Alternatively, pharmacological inhibitors can be used to reconstitute ISC niche signaling pathways *in vitro*. Indeed, organoids can be cultured with the GSK3 inhibitor CHIR-99021 (activating the Wnt pathway) and the BMP type I receptor inhibitor LDN-193189 (inhibiting the BMP pathway) (Li et al., 2018b).

Recently, the methods used to grow human or mouse intestinal organoids were successfully adapted to several other animal species (pig, cow, sheep, chicken, horse and dog) (Derricott et al., 2019; Hamilton et al., 2018; van der Hee et al., 2018; Powell and Behnke, 2017; Yin et al., 2019). However, a method to grow rabbit intestinal organoids is lacking despite the usefulness of such model in the context of biomedical or veterinary research (e.g. study of human bacillary dysentery or of epizootic rabbit enteropathy) (Licois et al., 2005; Yum et al., 2019). Herein, we present for the first time a method to grow rabbit caecum organoids in 3D. Our results show that replicating the ISC niche with pharmacological inhibitors or with L-WRN CM differentially regulates epithelial proliferation, differentiation and barrier function. We also show that rabbit caecum organoid cells can be cultured in 2D monolayers to facilitate the access to the apical side of IEC, which is needed to study their interactions with luminal components such as microbes, nutrients or xenobiotics.

## Materials and methods

### L-WRN cell culture

L-WRN cells (ATCC, Cat# CRL-3276 Manassas,VA) (passages 1-11) were seeded in T75 flasks at 37°C under 5% CO_2_ atmosphere in “complete DMEM” containing: Dulbecco’s Modified Eagle Medium (DMEM) high glucose GlutaMAX™ Supplement pyruvate (Thermo Fisher Scientific, Waltham, MA), 10% fetal bovine serum (FBS) (Thermo Fisher Scientific) and 1% penicillin/streptomycin (P/S) (Sigma-Aldrich, St-Louis, MO). Cells containing plasmids for expression of mouse Wnt3a, R-spondin 3 and Noggin were selected during the 3 first days post seeding by adding 0.5 mg/mL G418 (Sigma-Aldrich) and 0.5 mg/mL Hygromycin B (InvivoGen, San Diego, USA) to the complete DMEM (Miyoshi and Stappenbeck, 2013) (figure 1A). Then, cells were washed with PBS to remove residual antibiotics and cultured in complete DMEM without G418 nor Hygromycin B. The conditioned medium (L-WRN CM) was collected every day for 4 days. After centrifugation (24 g, room temperature, 5 min), L-WRN CM were kept at -20°C before pooling when 500 mL were produced. L-WRN cells were weekly sub-cultured (dilution 1:10 v/v) by partial digestion with EDTA-trypsin 0.25% w/v (Thermo Fisher Scientific).

**Figure 1:**
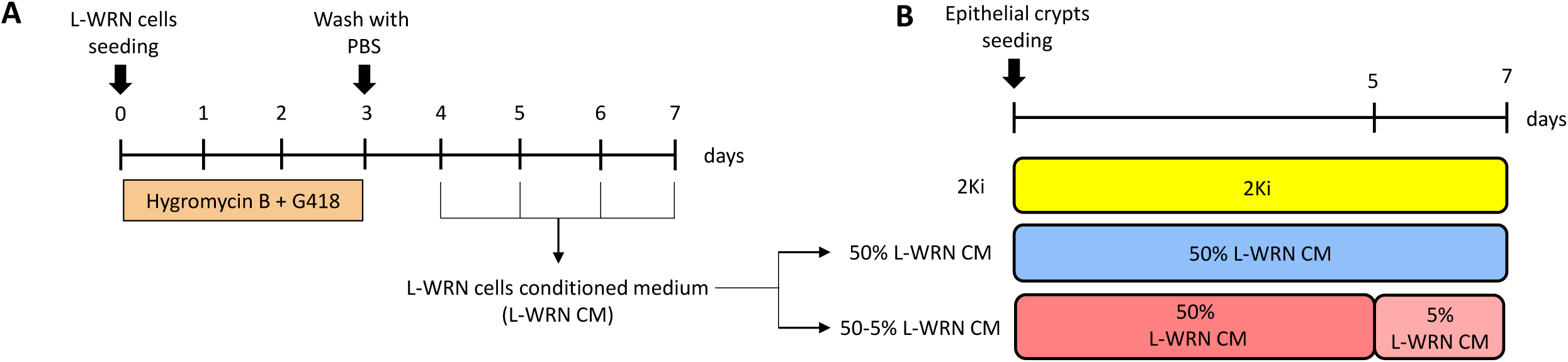
Schematic representation of methods used for organoid culture. **A.** L-WRN cells were mseeded in complete DMEM (DMEM, 10% SVF, 1% P/S) with Hygromycin B and G418 to select cells containing expression plasmids for mouse Wnt3a, R-spondin 3 and noggin. After 3 days of culture, cells were washed with PBS and cultured with complete DMEM only. Conditioned medium (L-WRN CM) was collected every 24 hours. **B.** Organoids were cultured for 7 days in 3 different conditions: (i) with pharmacological inhibitors (2Ki) for 7 days or (ii) with 50% L-WRN CM for 7 days or (iii) with 50% L-WRN CM for 5 days and then 5% L-WRN CM for 2 days.

### Isolation of intestinal crypts

Rabbits (line INRAE 1777) were bred at the PECTOUL Experimental Unit (INRAE, Castanet-Tolosan, France). Animals were handled according to the European Union recommendations on the protection of animals used for scientific purpose (2010/63/EU) and in agreement with French legislation (NOR:AGRG1238753A 2013). Animal experiments received the approval of the local ethical committee (SSA_2018_010). Caecal tissues used in this study were obtained from 30-day-old male rabbits slaughtered by electronarcosis and exsanguination. Epithelial crypts were isolated from rabbit caecal tissue fragments by incubation in a dissociation solution (PBS without Ca^2+^-Mg^2+^ (Thermo Fisher Scientific), 9 mM EDTA (Thermo Fisher Scientific), 3 mM 1,4-Dithiothréitol (DTT) (Roche, Basel, Switzerland), 10 µM ROCK inhibitor Y27632 (ATCC® ACS-3030™) and 1% P/S (Sigma Aldrich)) under agitation during 30 min at room temperature. Then, crypts were detached by manual shaking (1 min) in cold PBS. After centrifugation (24g, 4°C, 5 min), the crypt pellet was re-suspended in cold DMEM and used immediately for organoid culture.

### Organoid culture

Crypts were counted and re-suspended in ice-cold Matrigel (Corning, Corning, NY) and seeded in pre-warmed 48-well plates at a density of 150 crypts/25 µL of Matrigel. After polymerization (37°C, 5% CO_2_, 20 min), 250 µL warm organoid culture media (3 conditions detailed in table 1) was added as indicated in figure 1B. The media was replaced every 2-3 days. After 7 days of culture, organoids were counted according to their spherical or non-spherical morphology by bright field optical microscopy (Eclipse Ts2, Nikon Instruments, Melville, NY). The diameter of organoids and epithelial cell height were measured with Image J software (Schneider et al., 2012).

**Table 1:**
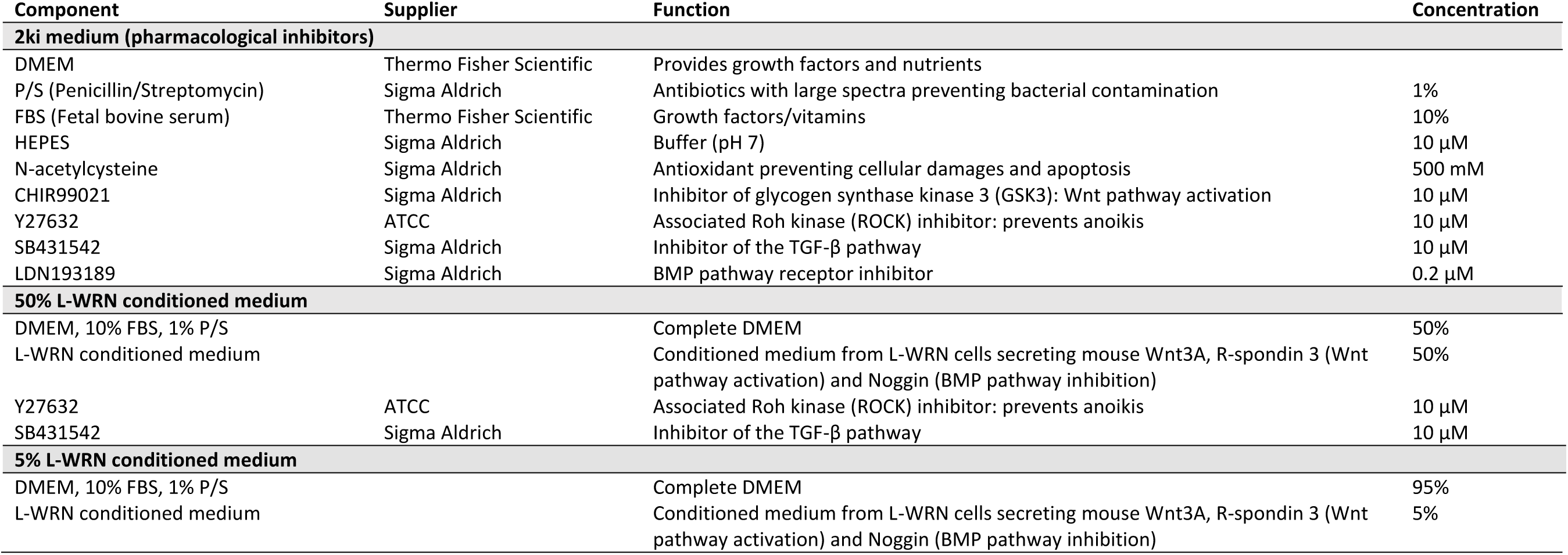
Composition of 2Ki and 50% or 5% L-WRN CM medium. L-WRN CM: conditioned medium produced by L-WRN cells

Every week, after washing with warm PBS, organoids were broken by pipetting in EDTA-trypsin 0.25% w/v (Thermo Fisher Scientifc) before incubation (37°C, 5% CO_2_, 5 min). Six wells were pooled before centrifugation (24g, 4°C, 5 min) and the pellet was re-seeded in Matrigel with a dilution ratio 1:8 and cultured as previously described. For organoids cryoconservation, the media was removed and freezing solution was added (80% complete DMEM, 10% FBS, 10% DMSO, 10 µM Y27632). Matrigel domes were scratched with a pipet tip and the pool of 6 wells were transferred in a cryotube and then placed in an isopropanol freezer box, allowing the cells to be gradually frozen at -80°C, before being kept in liquid nitrogen for long term storage. After fast thawing at 37°C, organoids were re-cultured with a dilution ratio 1:8 as described above.

### Two-dimensional monolayers of organoid cells

Organoids cultured in 3D with 2Ki medium or 50% L-WRN CM for 1 week were collected from a pool of 6 wells (about 1000 organoids) in complete DMEM before centrifugation (95 g, room temperature, 5 min). Organoids were dissociated into single cells by incubation in pre-warmed TrypLE Express Enzyme (Thermo Fisher Scientific) for 10 minutes at 37°C. After centrifugation (95 g, room temperature, 5 min), dissociated cells were resuspended in organoid culture medium and seeded in one Transwell (0.33 cm^2^ culture area) (Thermo Fisher Scientific). Transwells were previously coated by incubation (37°C, 5% CO_2_, 1 h) with diluted Matrigel (1:50 v/v in DMEM) before drying (room temperature, 10 min). 2Ki medium or 50% L-WRN CM was added at the apical (400 µL) and basal (600 µL) side. After 24h of incubation, the medium was replaced by complete DMEM for 24 h to allow IEC differentiation, as previously described (Moon et al., 2014; VanDussen et al., 2015).

### Gene expression

Organoids were washed with PBS and 6 wells/condition were pooled in 300 µL of TRI Reagent (Zymo Research, Irvine, CA) and kept at -80°C until total RNA extraction. After thawing, samples were centrifuged (12 000 g, 4°C, 10 min) to remove particles. The supernatant was collected and total RNA were extracted with Direct-zol RNA MiniPrep Plus kit (Zymo Research) according to the manufacturer’s instructions. A DNAse I digestion step was included to remove genomic DNA. After elution in RNase-free water, RNA concentration was determined using a Nanodrop before storage at -20°C until use. cDNA were prepared from 1 µg RNA with GoScript Reverse Transcription Mix, Random primer (Promega, Madison, Wisconsin) following the manufacturer’s instructions. cDNA were diluted 1:2 (v/v) and stored at -20°C. Gene expression was analyzed by real-time qPCR using QuantStudio 6 Flex Real-Time PCR System (Thermo Fisher Scientific) or Biomark microfluidic system using a 48.48 Dynamic Array IFC for gene expression (Fluidigm, San Francisco, CA) according to the manufacturer recommendations. The sequences of the primers used are presented in table S1. Data were analyzed with the 2^-ΔΔCt^ method with *Atp5b* gene expression used as a reference.

### Staining and confocal microscopy of organoids

Organoids were cultured as described above in 3D in Nunc Lab-Tek Chamber Slide system (Thermo Fisher Scientific). For proliferative cells staining, organoids (after 1 week of culture) were incubated (37°C, 5% CO_2_, 1 h) with 10 µM EdU (5-ethynyl-2’-deoxyuridine, Thermo Fisher Scientific), a nucleoside analog of thymidine incorporated into DNA during active DNA synthesis. After fixation with 4% paraformaldehyde (Electron Microscopy Sciences, Hatfield, PA) during 20 min under agitation, organoid cells were permeabilized with 0.5% triton X-100 (Sigma Aldrich) during 20 min under agitation. Between each step, wells were washed twice with 3% bovine serum albumin (Euromedex, Souffelweyersheim, France) in PBS. EdU was detected with Alexa 488 fluorochrome according to the protocol of Click-iT EdU Cell Proliferation kit (Thermo Fisher Scientific). For actin staining, organoids were fixed and permeabilized as described above before incubation with 10 µM phalloidin coupled with TRITC fluorochrome (Sigma Aldrich), under agitation (room temperature, 30 min). After staining, culture chambers were removed and mounting medium (Vector Lab, Burlingame, CA) supplemented with DAPI was added. Slides were stored in the dark at 4°C until observation. For organoid cell monolayers cultured in 2D, fixation and staining was performed as described above directly in Transwells before mounting the membrane on a microscope slide. Fluorescence staining was analyzed with a confocal laser scanning microscope (TCS SP8; Leica, Germany) using a x10 dry immersion objective lens or a x25 water immersion objective lens. DAPI, Alexa 488 and TRITC fluorescence was excited with the 405, 488 and 552 nm laser line and recorded in one of the confocal channels in the 415-475 / 500-550/ 560-620nm emission range, respectively. Images were acquired in the sequential mode using LAS X software (version 3.0, Leica).

### Protein sequence alignment

Mouse and rabbit orthologs of Wnt3a (NP_033548.1, XP_002723899.1), R-spondin 3 (NP_082627.3, XP_002714851.1) and Noggin (NP_032737.1, XP_002719325.1) were aligned with EMBOSS Needle (EMBL-EBI). The overall percentage of identity between mouse (GRCm38.p6) and rabbit (OryCun2.0) orthologous genes (ortholog_one2one, ortholog_one2many, ortholog_many2many) were obtained using the Biomart tool of the ENSEMBL database (Cunningham et al., 2019).

### Statistics

All statistical analysis were performed using the R software (version 3.5.1) with the packages car, lme4 and emmeans (version 3.5.1). A linear model was used to test the effect of each component of the 2Ki medium on organoid number or diameter. Mean values of each group (i.e. each composition of culture medium) were then compared to the mean value of the complete 2ki condition and P-values were adjusted with the Tukey method. In experiments comparing characteristics of organoids cultured with 2ki medium or L-WRN CM (50% or 50-5%), all conditions were tested on organoids from 4 rabbits and several conditions were tested on each culture plate. Thus a linear mixed model was used with medium composition as a fixed effect and rabbit and culture plate as random effects. Mean values of each group were compared pairwise and P-values were adjusted with the Tukey method. P<0.05 was considered significant.

## Results

### Reconstitution of the ISC niche with pharmacological inhibitors produces spherical rabbit caecum organoids

Since purified recombinant proteins (Wnt3a, R-spondin or Noggin) usually used to reconstitute the epithelial stem cell niche are not commercially available in rabbits, we first tested an organoid culture method called “2Ki” replacing these proteins by pharmacological inhibitors (Li et al., 2018b). After 7 days of culture in the 2Ki medium, spherical rabbit caecum organoids were obtained with a mean diameter of about 300 µm (figure 2 A-C). These organoids could be sub-cultured, cryopreserved and resuscitated (data not shown). Then, we removed one by one each component of the 2Ki medium to analyze their effects on the formation and growth of rabbit caecum organoids. FBS presence in the media was an absolute requirement to observe the formation of organoids. When Y27632 (ROCK inhibitor preventing IEC anoikis (Sato et al., 2009)) was removed, the number and the diameter of organoids tended to be lower than in complete 2Ki medium, although these changes were not statistically significant. In absence of CHIR-99021 (inhibitor of GSK3, activating of the Wnt pathway), only few organoids were observed and their diameter was very small (about 40 µM, 7-fold smaller than organoids cultured with the complete 2Ki medium). Removal of the TGF-β/BMP pathway inhibitors (SB-431542 and LDN-193189) did not modify significantly the number of organoids but reduced their diameter (1.5-fold smaller than organoids cultured with the complete 2Ki medium). Altogether, our results show that FBS and pharmacological inhibitors reproducing the stem cell niche (high Wnt/low BMP signaling pathway) are necessary for optimal growth of rabbit caecum organoids.

**Figure 2:**
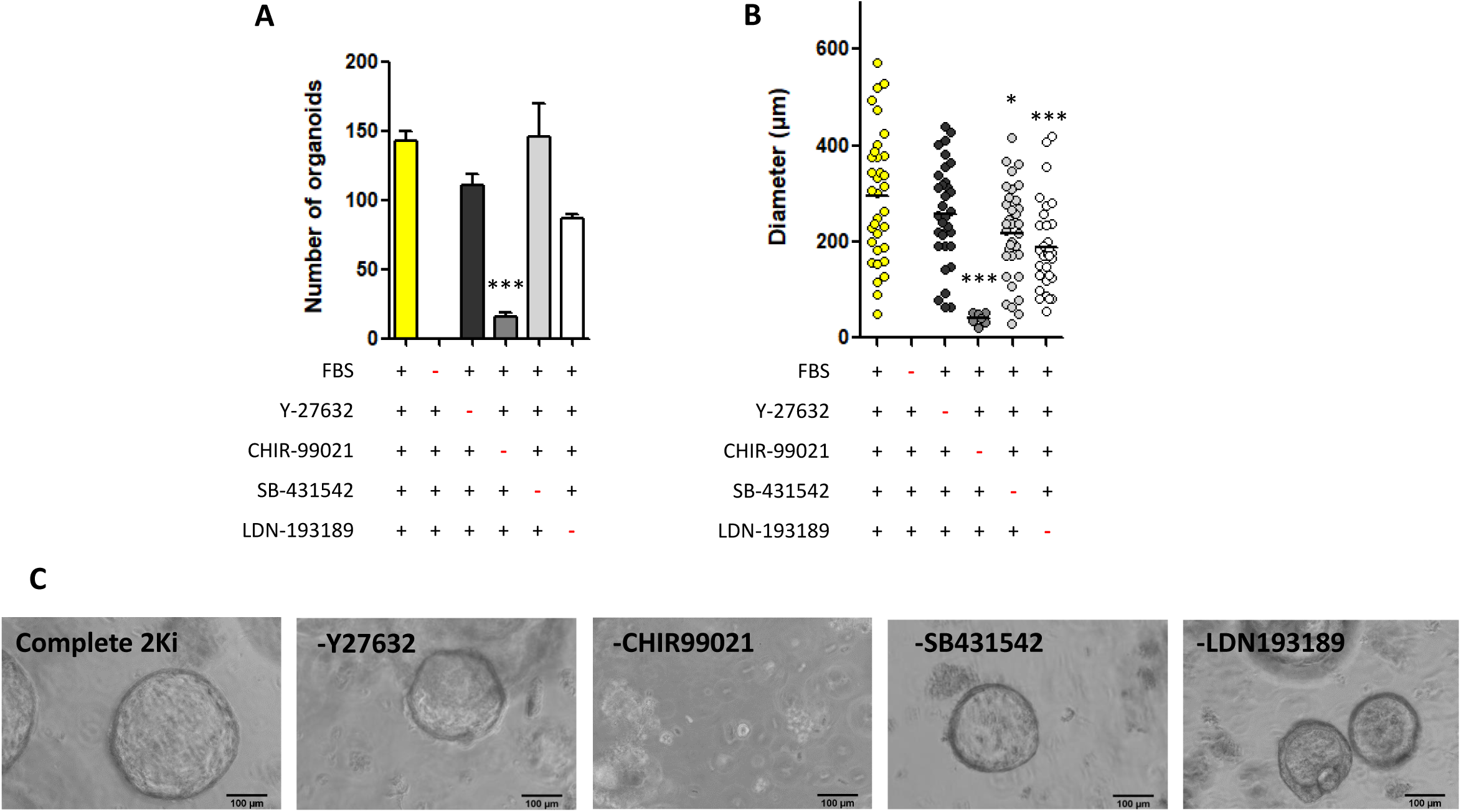
Characteristics of organoids cultured with pharmacological inhibitors (2Ki medium). Organoids were cultured during 7 days with complete 2Ki medium or without one of its components. **A.** Number of organoids per well (n=3). Data are presented as means + SEM. **B.** Diameter of organoids. Individual values are presented with dots and means with bars. A linear model was used to test the effect of the medium composition. Mean value of each group was compared to the mean value of the complete 2Ki condition. ***: P< 0.001, *: P < 0.05 (adjusted with the Tukey method). **C.** Representative images of organoids in each condition. The scale bar represents 100 µm.

### Reconstitution of the ISC niche with L-WRN CM produces rabbit caecum organoids with spherical and non-spherical morphologies

The spherical morphology of rabbit caecum organoids cultured with pharmacological inhibitors (2Ki medium) (figure 2C), suggested highly proliferative IEC with a low differentiation level (Merker et al., 2016). Such poorly differentiated culture might not be relevant to test interactions with nutrients or microbes. Thus, in an attempt to grow more differentiated organoids, we developed a second culture method to reconstitute the ISC niche by using the CM of L-WRN cells (L-WRN CM) secreting mouse proteins Wnt3a, R-spondin 3 (Wnt signaling potentiator) and Noggin (inhibitor of the TGF-β superfamily, inhibiting the BMP pathway) (Miyoshi and Stappenbeck, 2013). The amino acid sequence is highly similar between mouse and rabbit orthologs of Wnt3a (94%), R-spondin 3 (93.9%) and Noggin (98.3%) (figure S1, knowing that the overall percentage of identity between mouse and rabbit orthologous genes is ∼ 86%). This high level of conservation suggested that L-WRN CM could support the growth of rabbit caecum organoids, as previously observed for several other animal species (Powell and Behnke, 2017). We used the L-WRN CM at two concentrations: 50% for 7 days (50% L-WRN CM condition) or 50% for 5 days followed by 5% for 2 days (50-5% L-WRN CM condition) (figure 1B) based on a previous publication (VanDussen et al., 2015). When compared to the 2Ki medium, slightly more organoids were obtained in the 50% L-WRN CM condition but not with the 50-5% L-WRN CM condition (figure 3A). In contrast, the diameter of organoids cultured in L-WRN CM (50% or 50-5%) was significantly smaller than in organoids cultured with the 2Ki medium (figure 3B). L-WRN CM produced organoids with more diverse morphologies than the 2Ki medium since both spherical and non-spherical organoids (i.e. mulitlobulated, ∼12%) were observed with L-WRN CM (50% or 50-5%) (figure 3C and E). Moreover, the height of organoid cells was significantly greater with L-WRN CM (50% or 50-5%) compared to organoids cultured with 2Ki medium (figure 3D and E). Of note, there was no significant differences in the morphological features of organoids cultured with L-WRN CM used at 50% or 50%-5%. Overall, our results indicate that mimicking the ISC niche with L-WRN CM (containing mouse Wnt3a, R-spondin 3 and Noggin) allowed the growth of rabbit caecum organoids with spherical and non-spherical morphologies.

**Figure 3:**
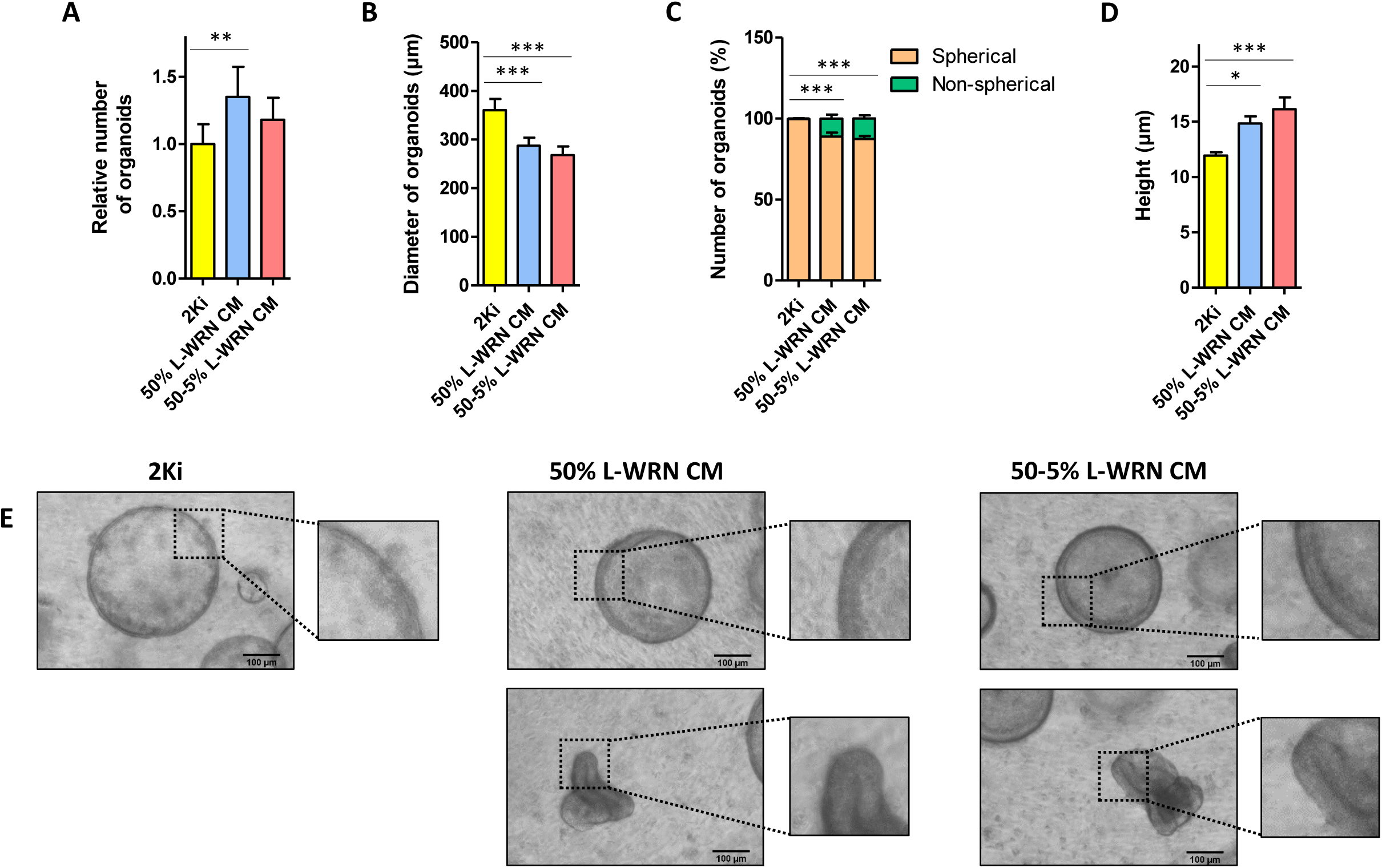
Comparison of characteristics of organoids cultured with 2Ki medium or L-WRN CM. Organoids were cultured during 7 days in 2Ki, 50% L-WRN CM (conditioned medium produced by L-WRN cells) or 50-5% L-WRN CM conditions. **A.** Number of organoids per well (expressed relatively to the number of organoids obtained for the same rabbit with 2Ki medium). **B.** Diameter of organoids. **C.** Number of organoids according to their spherical or non-spherical morphology. **D.** Cell height of organoids. **A-D.** Data are presented as means + SEM, n=7. A linear mixed model was used with medium composition as a fixed effect and rabbit and culture plate as random effects. Mean value of each group were compared pairwise. ***: P< 0.001, **: P<0.01, *: P < 0.05 (adjusted with the Tukey method). **E.** Representative images showing organoids in each condition. The scale bar represent 100 µm. Insets show a zoom to illustrate cell height.

### L-WRN CM induces less proliferation and more differentiation in rabbit caecum organoids than pharmacological inhibitors

The presence of non-spherical structures, the increased thickness of epithelial layer and the smaller diameter of organoids cultured with L-WRN CM suggested a lower proliferation rate and a higher differentiation level than in organoids cultured with pharmacological inhibitors (2Ki) (Merker et al., 2016). Therefore, to verify this hypothesis, we analyzed proliferation and differentiation markers in organoids grown in 2Ki medium or L-WRN CM. EdU staining suggested that less organoids were positive for proliferation in the 50% and 50-5% L-WRN CM conditions, when compared to 2Ki condition (figure 4A). However, in proliferating organoids (EdU^+^), the staining intensity was not different between the three conditions of growth media. Of note, we did not observe EdU positive cells in non-spherical organoids (L-WRN CM condition). The mRNA level of the stem cell marker *Lgr5* was the highest in organoids grown with 2Ki medium, while the expression of this gene was not detected in 1/4 samples with 50% L-WRN CM and 3/4 samples with 50-5% L-WRN CM (figure 4B). Similar trends were obtained for the other stem cell markers *Olfm4, Sox9* and *Smoc2*. In addition, the mRNA level of the proliferation marker *Pcna* was significantly reduced in organoids grown in 50-5% L-WRN CM, when compared to the 2Ki medium and 50% L-WRN CM (figure 4B). In contrast, markers of differentiated epithelial cells were highly expressed in organoids grown in 50-5% L-WRN CM, when compared to the 2Ki medium and, to a lower extent in 50% L-WRN CM (figure 5A and B). The gene expression of the absorptive cells markers *Krt20, Alpi, Vil1* and *Aqp8* were significantly higher in organoids grown in 50-5% L-WRN CM, when compared to the 2ki medium (figure 5A). *Vil1* was also significantly more expressed when L-WRN CM was used at 50-5% compared to 50%. The transcription factor involved in secretory lineage commitment *Atoh1* was significantly more expressed in organoids grown in L-WRN CM than in 2Ki medium (figure 5B). Goblet cells markers (*Muc2* and *Spdef)* and the enteroendocrine cells associated gene *Pyy* were more expressed in 50-5% L-WRN CM organoids compared to the 2Ki condition (figure 5B). Altogether, our results show that rabbit caecum organoids grown with pharmacological inhibitors (2Ki medium) were highly proliferative and enriched in stem cells, while organoids grown in L-WRN CM were more differentiated, notably when the L-WRN CM was used at low concentration for two days.

**Figure 4:**
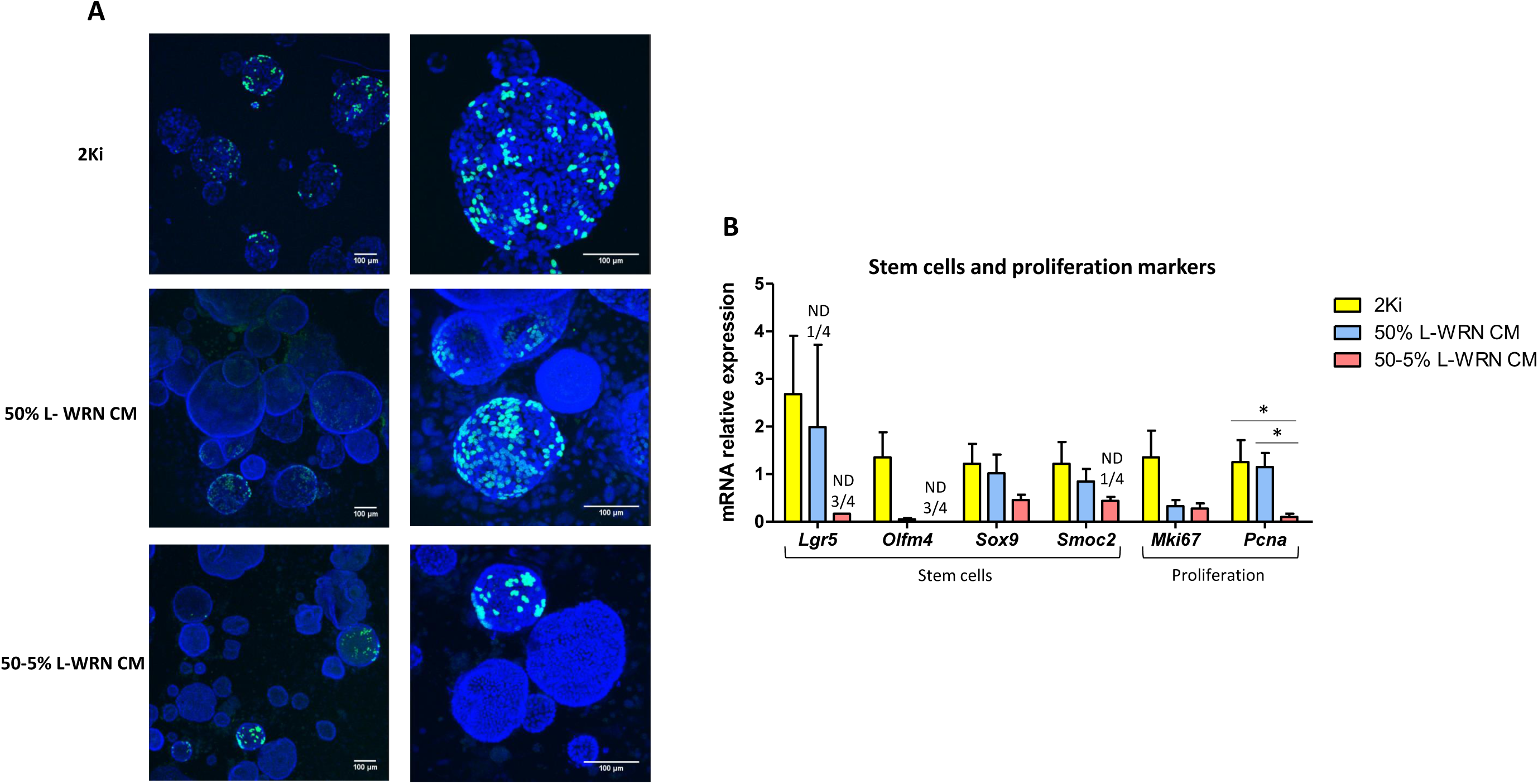
Comparison of proliferation markers in organoids cultured with 2Ki medium or L-WRN CM. Organoids were cultured during 7 days in 2Ki, 50% L-WRN CM (conditioned medium produced by L-WRN cells) or 50-5% L-WRN CM conditions. **A.** Proliferating cells were stained for 1 hour by EdU (green) in organoids after 7 days of culture. DAPI staining (blue) shows nuclei. The scale bar represents 100 µm. **B.** Relative gene expression of stem cells and proliferation markers. Gene expression was analyzed by high throughput microfluidic qPCR. Data are presented as means + SEM, n=4. A linear mixed model was used with medium composition as a fixed effect and rabbit and culture plate as random effects. Mean value of each group were compared pairwise. *: P < 0.05 (adjusted with the Tukey method). ND: non-detected.

**Figure 5:**
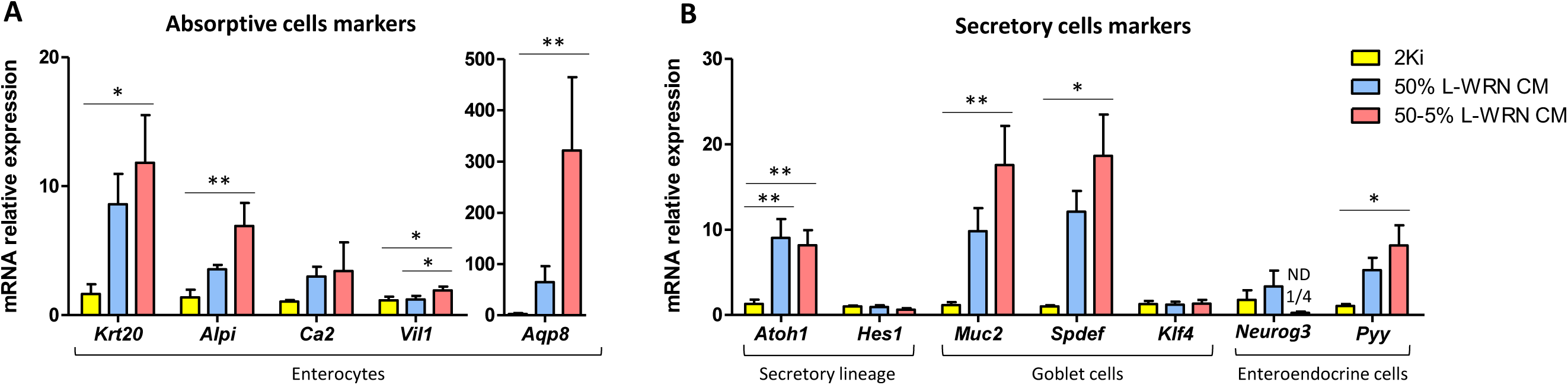
Comparison of differentiation markers in organoids cultured with 2Ki medium or L-WRN CM. Gene expression was analyzed by high throughput microfluidic qPCR in organoids cultured during 7 days in 2Ki, 50% L-WRN CM (conditioned medium produced by L-WRN cells) or 50-5% L-WRN CM conditions. **A-B.** Relative gene expression of absorptive cells markers (A) and secretory cells markers (B). Data are presented as means + SEM, n=4. A linear mixed model was used with medium composition as a fixed effect and rabbit and culture plate as random effects. Mean value of each group were compared pairwise. **: P<0.01, *: P < 0.05 (adjusted with the Tukey method). ND: non-detected

### Rabbit caecum organoids differentiation regulates the expression of key components of the epithelial barrier

As a next step, we explored whether the greater differentiation level observed in organoids cultured with L-WRN CM was associated with a modulation of the epithelial barrier function. The antimicrobial peptides *Reg3g* and *Defb1* were significantly more expressed in organoids grown in 50-5% L-WRN CM than in 2ki condition (figure 6A). Moreover, the expression level of *Reg3g* was also higher when the L-WRN CM was used at 50-5%, when compared to 50%. The mean expression of the antimicrobial peptide *Lyz* was more than 10-fold higher in organoids grown with L-WRN CM than in 2Ki medium, but this effect was not significant due to high variability. In contrast, the gene expression of the two monomers of the antimicrobial protein calprotectin (S100a8/S100a9) was strongly reduced or undetectable in organoids grown in L-WRN CM compared to 2Ki. The mRNA levels of most of the genes involved in epithelial junctions were not modified according to the culture condition excepted for *Cldn1* and *Cldn2* which mRNA levels was low or undetectable in organoids grow with L-WRN CM (figure 6B). Regarding genes involved in redox signaling in epithelial cells, there was a downregulation of the expression of *Gpx1* in organoids grown in L-WRN CM compared to the 2Ki medium (figure 6C). A similar effect was observed for the superoxide generating enzyme *Nox1* when L-WRN CM was used at 50%. In contrast, the nitric oxide producing enzyme *Nos2* was upregulated in organoids grown in 50% L-WRN CM compared to 2Ki and 50-5% L-WRN CM. The gene expression of the antioxidant enzyme *Sod1* was higher when the L-WRN CM was used at 50% compared to 50-5%. Finally, we observed no effect of organoid growth conditions on the mRNA levels of genes involved in cytokine signaling (*Il10ra*), toll-like receptor signaling (*Tlr2, Tlr5, Myd88*) or IgA secretion pathway (*Tnfsf13, Pigr*) (figure 6D). Altogether, our results show that the modulation of epithelial differentiation level in organoids induced by growth condition is associated with an alteration of the expression of several key components of the epithelial barrier (antimicrobial peptides, redox defenses and tight junctions).

**Figure 6:**
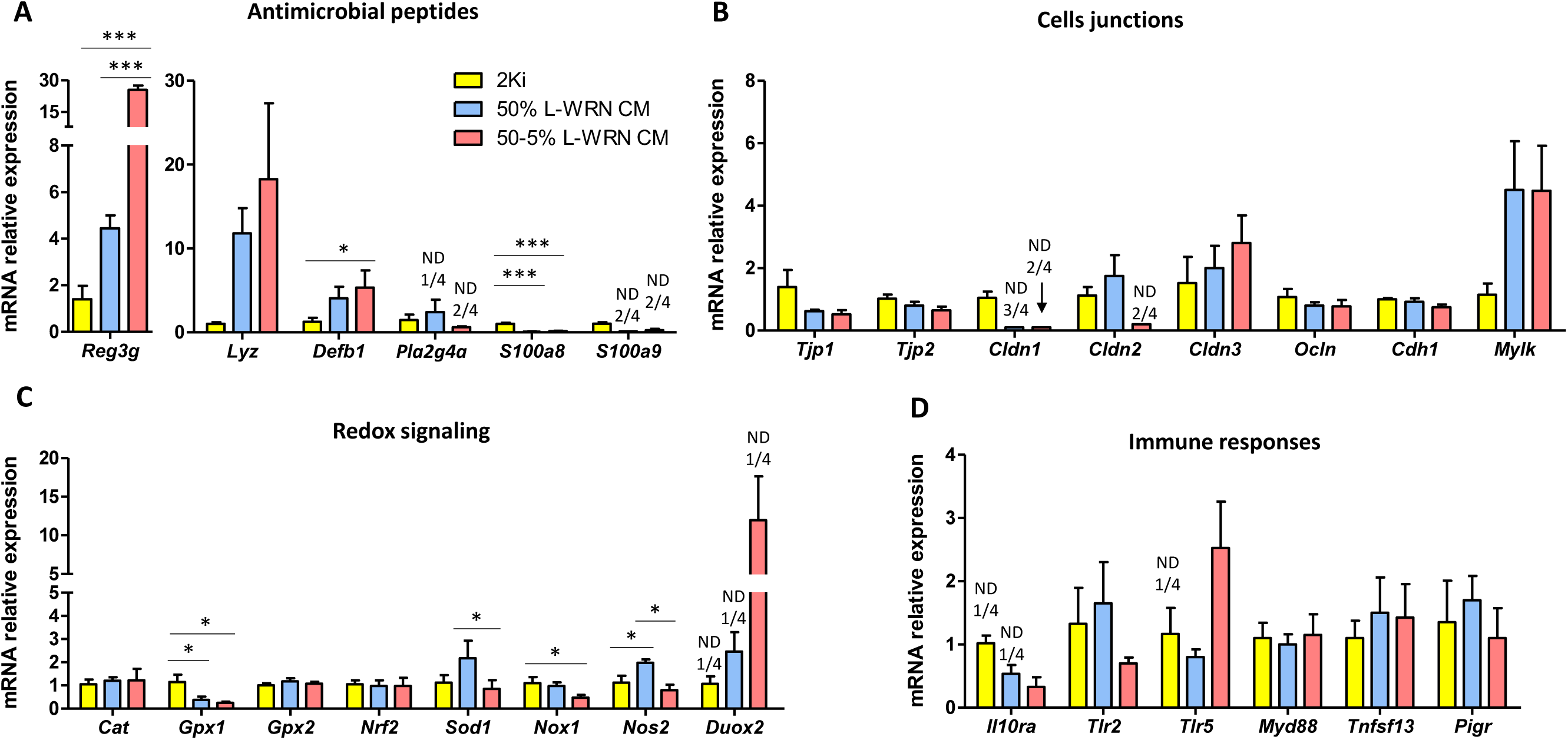
Comparison of epithelial barrier markers in organoids cultured with 2Ki medium or L-WRN CM. Gene expression was analyzed by high throughput microfluidic qPCR in organoids cultured during 7 days in 2Ki, 50% L-WRN CM (conditioned medium produced by L-WRN cells) or 50-5% L-WRN CM conditions. **A-D.** Relative expression of genes involved in antimicrobial defenses (A), cell junctions (B), redox signaling (C) and immune response (D). Data are presented as means + SEM, n=4. A linear mixed model was used with medium composition as a fixed effect and rabbit and culture plate as random effects. Mean value of each group were compared pairwise. ***: P<0.001, *: P < 0.05 (adjusted with the Tukey method). ND: non-detected

### Culture of rabbit caecum organoid cells in 2D facilitates the access to the apical side of epithelial cells

Accessibility of the apical side of epithelial cells is an essential feature of intestinal organoids, especially to study interaction between luminal compounds (nutrients, microorganisms) and IEC. Thus, we analyzed the polarity of rabbit caecum organoids cultured with pharmacological inhibitors (2Ki) or with 50% or 50-5% L-WRN CM. Actin (a marker of the apical side of epithelial cells (Co et al., 2019)) was detected towards the lumen of the organoids when cultured in 3D, in all culture condition tested (figure 7A). We then aimed at developing a model in which the apical compartment of rabbit cultured epithelial cells would be accessible. Rabbit organoids grown in 3D for 7 days with 2Ki medium or 50% L-WRN CM were dissociated into single cells before seeding in Transwells previously coated with a thin layer of Matrigel. Two days later, we observed a contiguous monolayer of epithelial cells (figure 7B). Actin was located towards the upper compartment and thus accessible in IEC from organoids grown in 2Ki medium or 50% L-WRN CM (figure 7B, Z-section). Interestingly, actin was localized exclusively at cell junctions in IEC of organoids cultured with 50% L-WRN CM while actin stress fibers were observed with the 2Ki medium. In conclusion, the culture of rabbit caecum organoid cells in 2D allows the accessibility to the apical side of epithelial cells using both 2Ki and L-WRN CM culture media.

**Figure 7:**
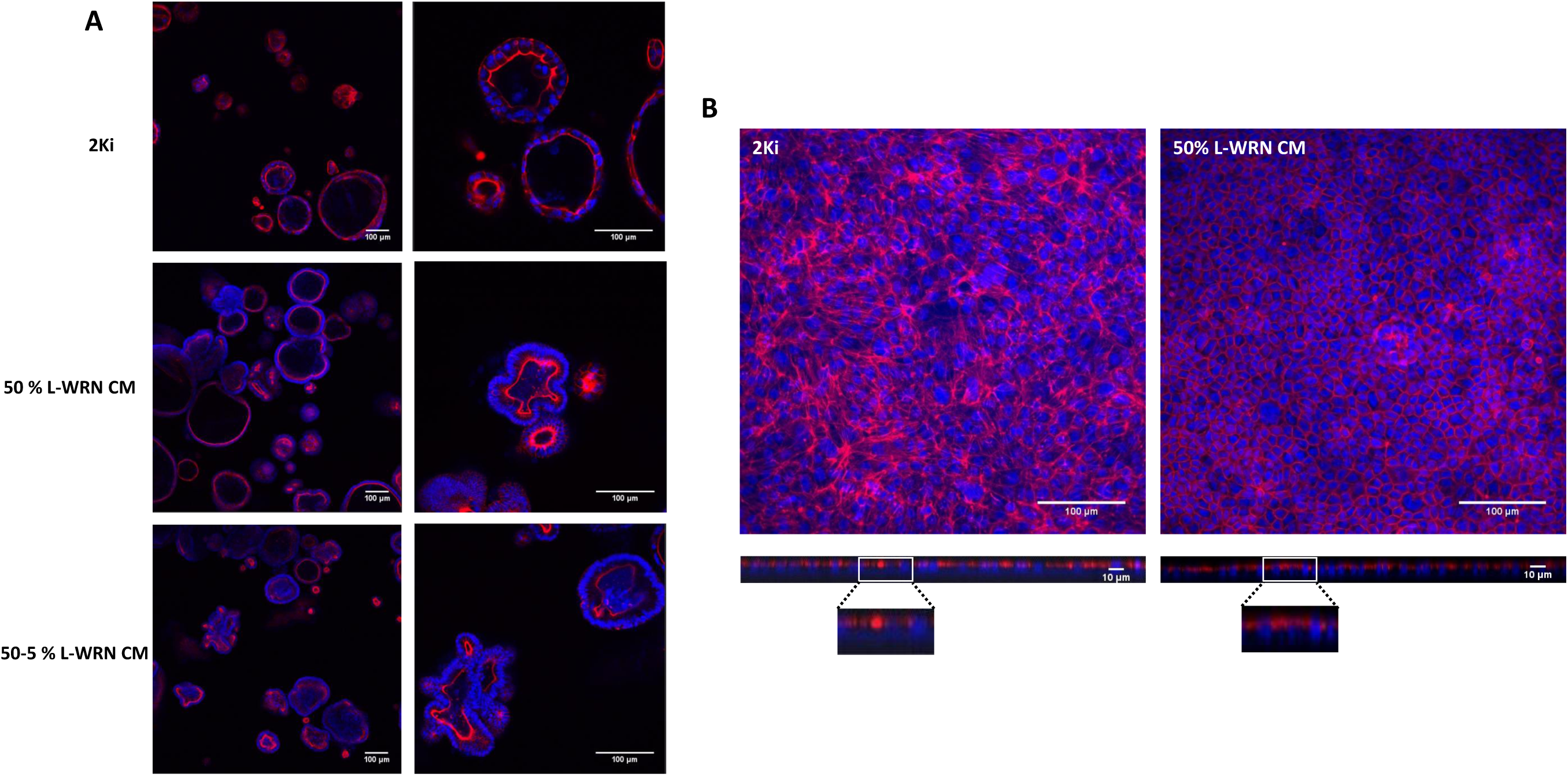
Polarization of organoids cultured with 2Ki medium or L-WRN CM. **A.** Organoids were cultured in 3 dimensions during 7 days in 2Ki, 50% L-WRN CM (conditioned medium produced by L-WRN cells) or 50-5% L-WRN CM condition. **B.** Organoids dissociated cells were cultured in 2 dimensions in 2Ki or 50% L-WRN CM conditions in insert coated with Matrigel. Phalloidin staining (red) shows actin and DAPI staining (blue) shows nuclei. The scale bar represent 100 µm (X-Y plan) or 10 µm (Z-section).

## Discussion

Our results show that rabbit caecum organoids can be obtained by seeding epithelial crypts in Matrigel with a growth culture medium reproducing *in vitro* the ISC niche either with pharmacological inhibitors or with L-WRN CM. Independently of the culture conditions, rabbit caecum organoids can be passaged, cryopreserved and resuscitated. These organoids contain the main epithelial cell types according to gene expression of markers for ISC and differentiated IEC (absorptive, goblet and enteroendocrine cells). Moreover, the gene expression of key components of the epithelial barrier (e.g. antimicrobial peptides, tight junction proteins, toll-like receptors) suggests that epithelial functionality is reproduced *in vitro*. Overall, the newly developed rabbit caecum organoid model closely replicates epithelial physiology and could thus be used to study rabbit epithelium in the context of biomedical or veterinary research.

To culture rabbit caecum organoids, we reconstituted the ISC niche *in vitro* either with pharmacological inhibitors or with L-WRN CM, based on previous publications (Li et al., 2018b; Miyoshi and Stappenbeck, 2013; Powell and Behnke, 2017; VanDussen et al., 2015, 2019). High Wnt/low BMP signaling pathways activation is needed for ISC maintenance and proliferation while low Wnt/high BMP signaling pathways activation leads to IEC differentiation (Gehart and Clevers, 2019). In our experiments, Wnt signaling pathway was activated by the GSK3 inhibitor CHIR-99021 in the 2Ki medium and by Wnt3a and R-Spondin 3 in L-WRN CM. BMP signaling was inhibited by LDN-193189 in the 2Ki medium and by Noggin in the L-WRN CM. Pharmacological inhibitors or mouse proteins secreted in L-WRN CM probably resulted in differential Wnt and BMP signaling pathway activation in rabbit caecum organoids which could explain the phenotypic differences observed according to the culture method. Moreover, the use of L-WNR CM at low concentration (5%) probably resulted in a low activation of Wnt signaling pathway and a low inhibition of the BMP pathway, as previously shown in human and mouse organoids (VanDussen et al., 2015, 2019).

The greater diameter of organoids cultured with 2ki medium, when compared to L-WRN CM indicated a faster epithelial growth with pharmacological inhibitors, suggesting a high Wnt/low BMP signaling pathways activation. Accordingly, removal of the Wnt activator CHIR-99021 or of the BMP inhibitor LDN-193189 from the 2Ki medium reduced the diameter of organoids and thus limited their growth. Organoids cultured with 2Ki medium were exclusively spherical/cystic, this spheroid morphology being associated with an immature phenotype and a high proliferation rate (Merker et al., 2016; Stelzner et al., 2012; VanDussen et al., 2019). In contrast, organoids cultured with L-WRN CM were not all spheroids, some of them being multilobulated, suggesting that these organoids were more mature, less proliferative than organoids cultured with 2Ki medium. Indeed, we did not observe EdU^+^ cells (i.e. proliferating) in non-spherical organoids cultured with L-WRN CM. In contrast, spherical organoids positive for EdU staining were observed both with 2Ki medium and L-WRN CM, but their frequency seemed to be higher with the 2Ki medium. Accordingly, the gene expression of the ISC markers *Lgr5* and *Olfm4* (Flier et al., 2009; Sato et al., 2009) and of the proliferation marker *Pcna* was very low or not detectable when the L-WRN CM was used at low concentration. Our results are in agreement with a previous study in human organoids showing that reducing the L-WRN CM from 50% to 5% downregulated the expression of *Lgr5* and *Olfm4* (VanDussen et al., 2015). In another study, mouse colon organoids cultured with 2Ki medium had higher expression of *Lgr5* than organoid cultured with a medium containing recombinant Noggin and R-spondin (Li et al., 2018b), suggesting high Wnt/low BMP signaling pathways activation in organoids cultured with 2Ki medium. Besides Wnt and BMP signaling pathway, epidermal growth factor (EGF) is a key regulator of IEC proliferation and is an essential component of organoid culture media (Gehart and Clevers, 2019). Our two culture methods (2Ki and L-WRN CM) contained FBS which is a source of EGF (Miyoshi and Stappenbeck, 2013). This could explain why we did not obtain organoids when the 2Ki medium did not contain FBS. However, a study showed that EGF was not needed to grow mouse colon organoids with the 2Ki medium containing the supplements N-2 and B-27 (Li et al., 2018b). Thus, in our culture conditions, FBS might bring essential but undefined compounds such as insulin or vitamins contained in N-2 and B-27 supplements. Altogether, our results show that the 2Ki medium is a more potent inducer of ISC maintenance and proliferation in rabbit caecum organoids than the L-WRN CM, especially when the latter is used at low concentrations.

Cell height was greater in organoids cultured with L-WRN CM than with 2Ki medium, this characteristic being typical of differentiated and highly polarized IEC (Altay et al., 2019). Accordingly, the gene expression of absorptive cells markers (*Krt20, Alpi, Aqp8* and *Vil1*) was higher in organoids cultured with L-WRN CM (notably when used at 50-5%) compared to the 2Ki medium. These results obtained in rabbit caecum organoids are consistent with previous data obtained in mouse organoids showing that the 2Ki medium reduced the gene expression of *Alpi* when compared to a growth medium containing recombinant EGF, Noggin and R-spondin (Li et al., 2018b). Upregulation of absorptive cells markers in human organoids were also found when L-WRN CM was used at low concentration (VanDussen et al., 2015). The gene expression of *Atoh1* coding for the transcription factor Math1, required for the commitment of epithelial progenitors into the secretory lineage (Yang et al., 2001), was upregulated in organoids cultured with L-WRN CM compared to the 2Ki medium, suggesting a higher number of secretory cells. This result was associated with an upregulation of goblet cells markers (*Muc2* and *Spdef*) and of the enteroendocrine cell gene *Pyy* in organoids cultured with L-WRN CM, especially when used at 50-5%. Altogether, our results indicated that epithelial cells were more differentiated when cultured with L-WRN CM used at low concentration compared to the 2Ki medium. IEC differentiation is usually associated with a modulation of epithelial barrier function (Sun et al., 2017). Indeed, in our experiments, differentiation of rabbit caecum organoid was associated with the regulation of antimicrobial peptides and proteins involved in cell junction and redox signaling. Overall, organoids cultured with 2Ki medium were highly proliferative and contained few differentiated cells while the opposite was observed for organoids cultured with L-WRN CM. These phenotypes are consistent with a higher activation of Wnt pathway and more potent inhibition of BMP signaling pathway in rabbit caecum organoids cultured with the 2Ki medium than with L-WRN CM.

When rabbit caecum organoids were cultured in 3D with 2Ki medium or L-WRN CM, the apical side of the epithelium was oriented toward the lumen of organoids, according to actin localization. Similar results were previously obtained in 3D intestinal organoids from human, mouse or farm animals (Co et al., 2019; Derricott et al., 2019). Thus, the physiological route of exposure to luminal compounds (microbes or nutrients) is not easily accessible in 3D rabbit caecum organoids. To circumvent this issue, we seeded dissociated cells from 3D organoids onto Matrigel-coated Transwell permeable supports, based on previous publications (van der Hee et al., 2018; Moon et al., 2014; VanDussen et al., 2015). We obtained 2D contiguous monolayers, in which IEC apical side was facing upwards, making it accessible to experimental treatments. Interestingly, the presence of actin stress fibers in 2D monolayers cultured with 2Ki medium but not with L-WRN CM is consistent with our results indicating a lower differentiation level when the 2Ki medium is used. Indeed, actin stress fibers are typically observed in IEC with a low differentiation level, while the perijunctionnal localization of actin (as observed with the L-WRN CM) is associated with a more differentiated state (Li et al., 2018a; Soubeyran et al., 1999).

In summary, reconstitution of the ISC niche with pharmacological inhibitors or with L-WRN CM allowed the growth of rabbit caecum organoids. Due to the large phenotypic differences observed according to the method used, we anticipate different applications of organoids cultured with 2Ki medium or L-WRN CM. Rabbit caecum organoids cultured with 2Ki medium could be used to generate rapidly a large number of rabbit ISC *in vitro.* In contrast, the L-WRN medium could be used to culture organoids composed mainly of differentiated IEC and thus appropriate to study rabbit epithelium physiology. Our results also show that intestinal organoid culture methods developed in mice or human can be easily adapted to other animal species, probably due to the high evolutionary conservation of the key regulators of ISC niche signaling pathways. Thus, we believe that the culture of intestinal organoids should be feasible for most animal species for which *in vitro* models of IEC are not available yet.

## AUTHOR CONTRIBUTION

EM, SC and MB: conceived the experiments. EM, CP, VH, SF and MB performed experiments. CC, AR and NV shared expertise and biological resources. EM, SC, GP and MB analyzed the data. EM, SC and MB wrote the manuscript. All authors contributed to manuscript editing.

## ACKNOWLEDGEMENTS

The authors thank the staff of the rabbit experimental unit PECTOUL for animal care. We also acknowledge Yvette Lahbib-Mansais and Christelle Marrauld (GenPhySE, INRAE) for help with cell culture and confocal imaging experiments.

## Abbreviations

2D: 2 dimensions
2Ki: name of the organoid culture medium containing pharmacological inhibitors only
3D: 3 dimensions
BMP: bone morphogenic protein
CM: conditioned medium
DMEM: Dulbecco’s Modified Eagle Medium
FBS: fetal bovine serum
IEC: intestinal epithelial cell
ISC: intestinal stem cell
L-WRN: L cell line engineered to secrete mouse Wnt3a, R-spondin 3 and Noggin
P/S: penicillin/streptomycin

## SUPPLEMENTAL FIGURE LEGENDS

**Figure S1:**
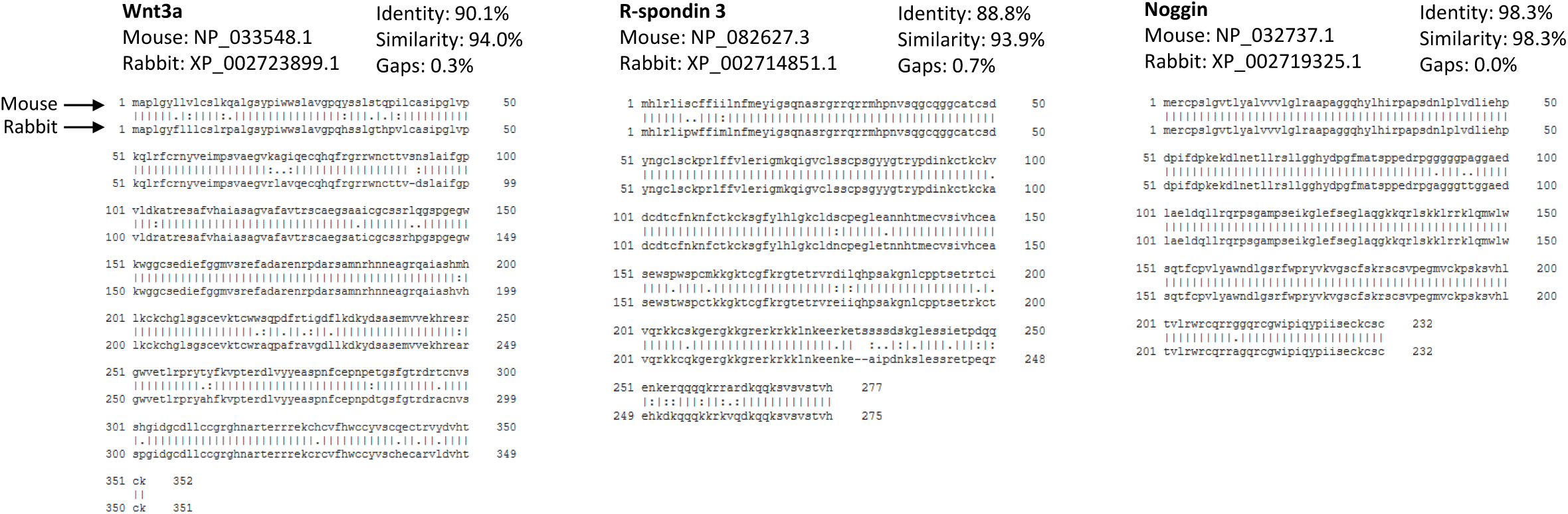
Amino acid sequence alignment of Wnt3A, R-spondin 3 and Noggin protein orthologs from mouse (upper row) and rabbit (lower row).

## SUPPLEMENTAL MATERIALS

**Table S1:**
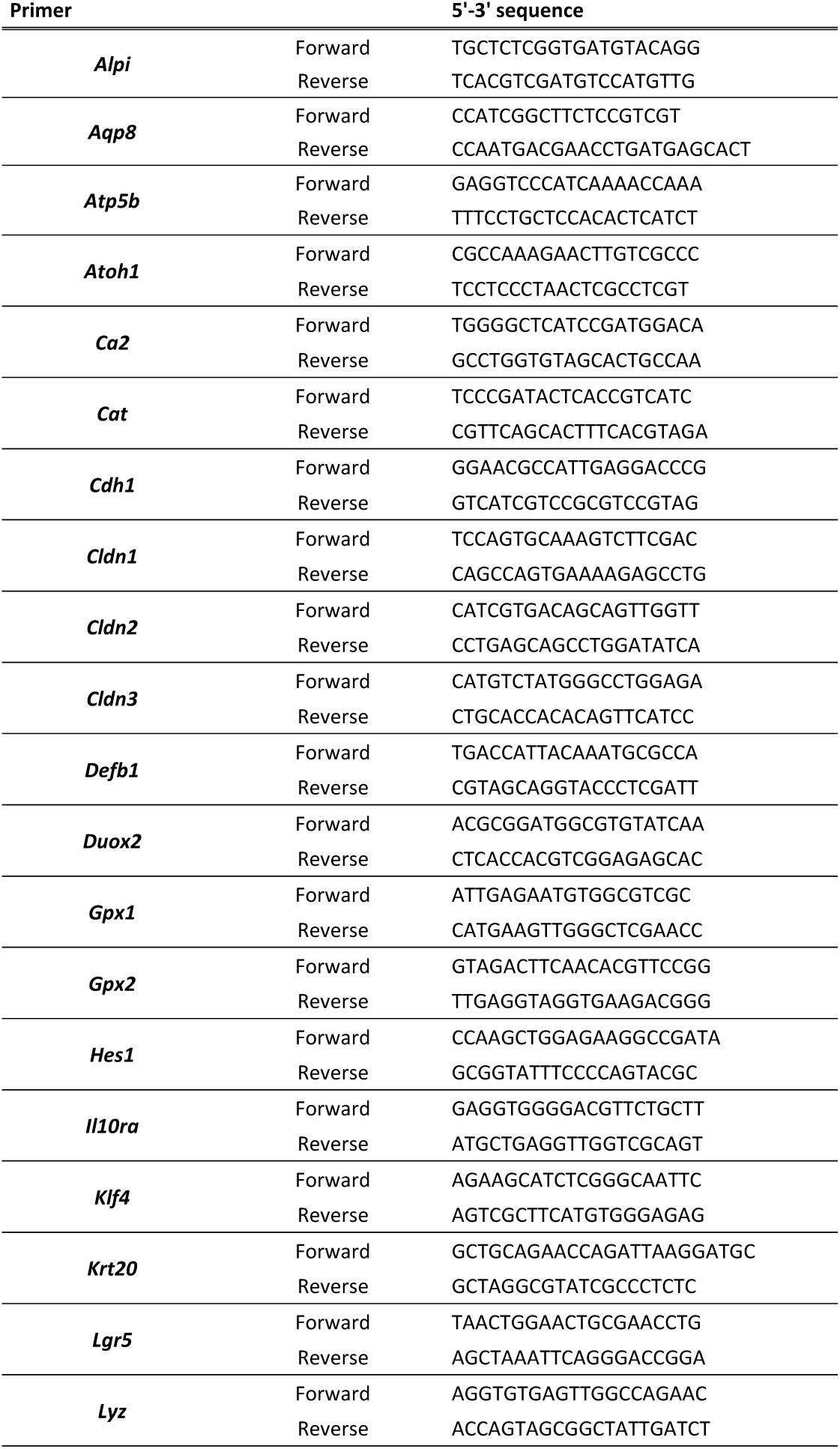
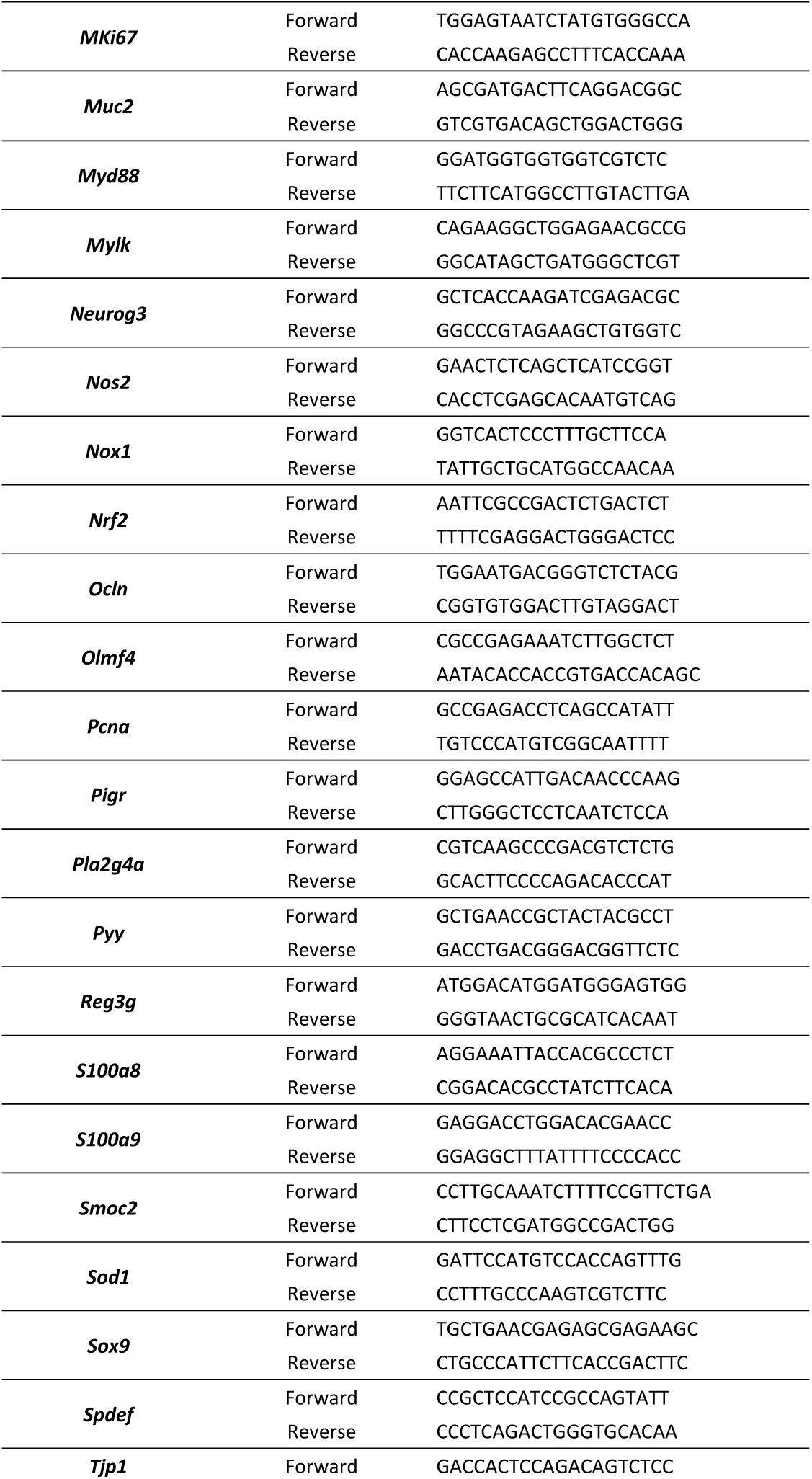
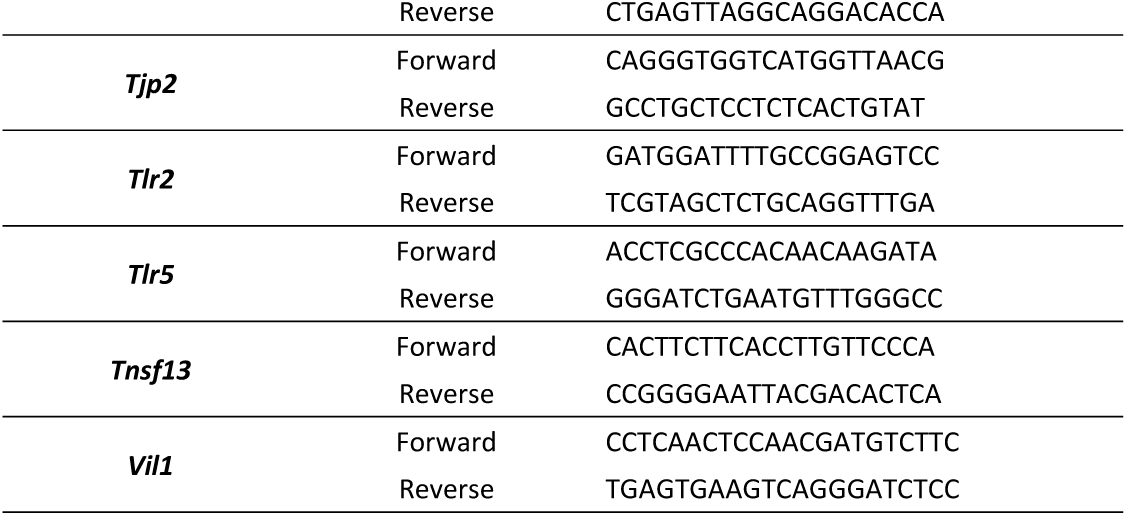
List of oligonucleotides used for qPCR

